# Risk of recurrent pregnancy loss in the Ukrainian population using a combined effect of genetic variants

**DOI:** 10.1101/603431

**Authors:** E. M Loizidou, A. Kucherenko, P. Tatarskyy, S. Chernushyn, G. Livshyts, R. Gulkovskyi, I. Vorobiova, Y. Antipkin, O. Gorodna, M. A. Kaakinen, I. Prokopenko, L. Livshits

## Abstract

Recurrent pregnancy loss (RPL) affects nearly 5% of the women of reproductive age. Its heterogeneous and multifactorial nature complicate both diagnosis and treatment, as well as identification of the genetic contribution to RPL. Evidence about the aetiology of RPL is controversial; however, several biological mechanisms have been proposed. Given the current knowledge about the genetic susceptibility to idiopathic RPL, we aimed to evaluate the predictive ability of a combined variant panel to the risk of RPL in the Ukrainian sample of 114 cases and 106 healthy controls. We genotyped variants within the 12 genetic loci reflecting the main biological pathways involved in pregnancy maintenance: blood coagulation (*F2, F5, F7, GP1A*), hormonal regulation (*ESR1, ADRB2*), endometrium and placental function (*ENOS, ACE*), folate metabolism (*MTHFR*) and inflammatory response (*IL6, IL8, IL10*). We showed that a genetic risk score (GRS) calculated from the 12 variants was associated with an increased risk of RPL (odds ratio 1.56, 95% CI: 1.21,2.04, *P=*8.7×10^−4^). The receiver operator characteristic (ROC) analysis resulted in the area under the curve (AUC) of 0.64 (95% CI: 0.57, 0.72), indicating an improved ability of the GRS to classify women with and without RPL. In summary, implementation of the GRS approach can help defining women at higher risk to complex multifactorial conditions such as RPL. Future well-powered genome-wide association studies will help in the dissection of biological pathways not hypothesised previously for RPL and further improve the prediction and identification of those at risk for RPL.

## Introduction

The loss of two or more sequential pregnancies in the first trimester of gestation is defined as recurrent pregnancy loss (RPL)(1). Nearly one in twenty women of reproductive age is affected by this condition(2). Heterogeneity and multifactorial nature of RPL complicate both diagnosis and treatment of RPL, thus causing severe distress for affected couples and their clinicians(3,4). Despite a large number of clinical and genetic studies aiming to identify probable causes and suitable treatments of RPL, most of the findings remain controversial and demand replication(3,5).

The vast majority of early pregnancy losses (50%–60%) are the consequence of chromosomal abnormalities, which can be of parental origin, or arise de novo in the embryo from parents with normal chromosomes(6,7). Nonetheless, endocrine, immunological, anatomical and other hypotheses are proposed to play a leading role in RPL aetiology, with most of the remaining RPL cases being idiopathic(5,8). In patients with idiopathic RPL, a multifactorial nature of the condition is usually suggested, with genetic component viewed as an important risk factor(5,9,10). Large-scale genome-wide association studies (GWAS) in appropriately defined individuals should enable better dissection of pregnancy maintenance/loss mechanisms, as well as provide evaluation of miscarriage risks in couples with reproductive complications. However, the genetic susceptibility to RPL in female health has not been addressed. The identification of RPL cases is laborious and expensive, hindering the setup of a well-powered GWAS for this outcome. Many studies, including the UK Biobank consisting of 500,000 individuals, have collected data on self-reported miscarriages and the number of spontaneous miscarriages. GWAS on the UK Biobank data(11) on such surrogate phenotypes for RPL have not resulted in any common variants associated with RPL at genome-wide significance, plausibly reflecting an inaccurate re-call and complex genetic susceptibility to RPL.

While more precise data on idiopathic RPL are being collected allowing well-powered GWAS in the future, our best approach is to look at the potential biological mechanisms proposed for RPL pathogenesis and use such information to infer the potentially relevant genes in RPL susceptibility. The suggested mechanisms include alterations in blood coagulation, hormonal regulation, endometrium and placental function, folate metabolism and inflammatory response.

### Blood coagulation

Haemostasis-related genes, such as the coagulation factor II and V genes *(F2* and *F5*, respectively), have been linked to venous thromboembolism and thrombosis in recent GWAS(12-15). They contribute to hereditary thrombophilia, and it has been suggested that they could act throughout pregnancy, causing miscarriages(16). Indeed, a study on *F2* and *F5* genes has shown associations with variation in these genes and RPL(17). Associations between polymorphisms in the coagulation factor VII (*F7)* gene and recurrent miscarriages were also reported in a study from Poland(18). Another strong candidate within the blood coagulation pathway to RPL is the glycoprotein Ib (platelet), alpha (*GP1A*) gene which is an important player in the platelet adhesion to collagen(19).

### Hormonal regulation

The relevance of hormonal regulation in the susceptibility of RPL comes from the knowledge that estrogens modulate multiple reproductive functions, including progesterone production and uteroplacental blood flow(20). Estrogen receptors (ERs), which in human consist of ERα and ERβ, encoded by *ESR1* and *ESR2* genes, respectively, are mediators of estrogen signalling and function (21,22). A study of *ESR1* locus variants and RPL did not yield support for the association(23); however, recent GWAS have linked variation within *ESR1* to susceptibility to endometriosis(24) and to maternal age at first birth(25). Stress-induced adrenergic receptor (ADRB2) activation may in turn directly affect embryo-maternal interactions during implantation, resulting in pregnancy complications and miscarriage (26). No studies have directly assessed the association with variation in this gene and RPL; however, controversial reports for the effects of *ADRB2* variants have been reported on preterm delivery (27,28) and its potential role was suggested as being a drug target for prevention of preterm delivery (29).

### Endometrium and placental function

Abnormalities of placental vasculature in the chorionic villi of RPL patients can be determined through low expression levels of angiogenesis-related genes, such as the endothelial nitric oxide synthase *(ENOS)* and angiotensin I-converting enzyme *(ACE)* genes. Variability within these genes may result in gestational complications, including pregnancy loss, pre-eclampsia, intrauterine foetal death and growth restriction (30). More specifically, variability in *ACE* is associated with RPL (31). The *ACE* gene, which generates angiotensin II from angiotensin I as a potent vasopressor (32,33), is a key component of the rennin–angiotensin system (RAS) that affects homeostasis (34). There is evidence that the presence of the D allele or D/D genotype is correlated with elevated plasma and tissue-specific ACE activity (35,36). In turn, eNOS is the main enzyme required for vascular NO production, by converting L-arginin to L-citrulline. *eNOS* is expressed in the terminal chorionic villous vessels and in the cyto- and syncytio-trophoblast layers during the first trimester (37).

### Folate metabolism

Variation within the 5,10-methylenetetrahydrofolate reductase (*MTHFR*) gene is associated with higher homocysteine (38,39) and serum folate concentrations (40). Mild elevations in the total plasma homocysteine (tHcy) concentration, a risk factor for placental abruption, and infarction and preeclampsia (41), are also associated with an increased risk of RPL (42–46). Furthermore, Nelen *et al*. demonstrated that the homozygosity for a common 677C→T mutation in the *MTHFR* gene leads to a two-fold to three-fold higher risk of RPL (46). Another association study between *MTHFR* and RPL was inconclusive, despite a large sample size of 1,830 cases and 3,037 controls (47).

### Inflammation

Cytokines form a complex regulatory network which maintains homeostasis between the foetus and maternal immune system. If this delicate balance is adversely affected, immunoregulatory mechanisms may be insufficient to restore homeostasis and this may lead to pregnancy failure (48). Variation especially at the inflammatory gene *IL10* has been associated with REPL (OR=3.01, P-value<10^−4^)(49), whereas the evidence between its association with RPL has been less conclusive (50), similarly to *IL6* and RPL (51). In addition, the chemokine IL-8 is a crucial player in the process of implantation, promoting trophoblastic cells migration and invasion (52). Moreover, microRNA studies for endometriosis have shown the potential role of IL-8 levels in the pathogenesis of endometriosis via stimulating endometrial stromal cell invasiveness (52).

Given the current knowledge about the genetic contribution to idiopathic RPL, we aimed to evaluate the predictive ability of a combined gene set of RPL associated DNA variants to the risk of RPL in a Ukrainian sample of 114 cases and 106 healthy controls. We genotyped variants at/within 12 genes reflecting the main biological pathways involved in pregnancy maintenance: blood coagulation (*F2, F5, F7, GP1A*), hormonal regulation (*ESR1, ADRB2*), endometrium and placental function (*ENOS, ACE*), folate metabolism (*MTHFR*) and inflammatory response (*IL6, IL8, IL10*).

## Material and Methods

### Study sample

REPLIK (REcurrent Pregnancy Loss In Kiev) study case group comprised 114 unrelated women at a mean age of 34.2 (SD 4.5) years with idiopathic RPL history undergoing observation in the State Institution “Institute of Paediatrics, Obstetrics and Gynaecology of NAMS of Ukraine” and perinatal clinic “ISIDA”. All the women were of Ukrainian territory descent from across Ukraine. RPL diagnosis was determined in case of at least two consequent miscarriages in the first trimester (mean number of foetal losses 2.7, SD 0.9). The American Society for Reproductive Medicine defines RPL as two or more clinical pregnancy losses, not necessary consecutive, documented by ultrasonography or histopathologic examination(53). In order to ensure the idiopathic nature of RPL in the studied patients, the following enrolment criteria were set: absence of a family history of birth defects; absence of the genital tract anatomic abnormalities, confirmed by ultrasonography or hysterosalpingography; a normal karyotype of both the studied individual as well as their partner, defined by GTG-banded chromosome analysis including GTG-banded metaphase plates with a minimum resolution of 400–450 bands per each sample. Moreover, blood tests for immunologic risk factors (anti-nuclear antibodies, anti-phospholipid antibodies, lupus anticoagulant), defects of thyroid function, diabetes mellitus, hyperprolactinemia and infections such as chlamydia were performed, and none of the individuals positive were included into the study group. A control group comprised 106 unrelated healthy women at a mean age of 26.2 (SD 3.0) years with no history of RPL or other pregnancy complications, no foetal losses, and have given birth to at least one naturally conceived child. Prior to clinical examination and genotyping, all participants had given their informed consent. The study has been approved by The Bioethical Committee of Institute of Molecular Biology and Genetics of NAS of Ukraine.

### Blood sample collection, DNA extraction and genotyping procedures

Venous blood samples from patients and control group individuals were collected into 4ml vacutainer tubes containing EDTA. Genomic DNA was extracted from blood samples by standard method using proteinase K with the following chlorophorm extraction. We used 13 genetic variants previously associated with RPL in European populations or those representing genetic networks of pathological processes leading to RPL (**Table 1** provides information about variants and reference papers). A SNP located in *IL10* gene (rs1800872) was excluded from the polygenic risk score calculation due to high LD (r^2^=0.26 in Europeans) with a nearby SNP included in the study (rs1800896). Genotyping for selected polymorphic variants was performed by common variations of PCR-based assays as described previously (**Table 1**) with slight modifications. We identified genotypes for all genotyped individuals without any missing values. However, genotyping of the 13 variants was done for the individuals depending on their time of joining the project and availability of the reagents (**Table 2** shows the numbers of cases and controls genotyped for each variant).

**Table 1.** Genetic variants used to assess their effect on the risk of RPL in the REPLIK study.

| Locus name | SNP rsID | EA/NE A | EAF (1000 G*) | OR (95% CI) | P-value | Case/Control sample size | Outcome | Reference | Genotyping method | Reference |
| --- | --- | --- | --- | --- | --- | --- | --- | --- | --- | --- |
| *Blood coagulation* |  |  |  |  |  |  |  |  |  |  |
| *F2* | rs1799963 | A/G | 0.008 | 2.00<br>(1.00,4.00) | < 0.03 | 342/123 | RPL | (17) | RFLP | (60) |
| *F5* | rs6025 | T/C | 0.99 | 2.5<br>(1.80,3.40) | < 10<sup>-3</sup> | 342/123 | RPL | (17) | RFLP | (60) |
| *F7* | rs6046 | G/A | 0.89 | -17 | - | - | - | - | RFLP | (61) |
| *GPIA* | rs1126643 | T/C | 0.40 | - | - | - | - | - | RFLP | (62) |
| *Hormonal regulation* |  |  |  |  |  |  |  |  |  |  |
| *ESR1* | rs2234693 | T/C | 0.58 | 1.10<br>(0.57,2.13) | > 0.05 | 350/646 | RPL | (56) | RFLP | (63) |
| *ADRB2* | rs1042714 | G/C | 0.41 | - | - | - | - | - | RFLP | (64) |
| *Endometrium and placental function* |  |  |  |  |  |  |  |  |  |  |
| *ENOS* | Intron-4†<br>VNTR | B/A | - | 1.005<br>(0.74,1.37) | > 0.05 | 410/357 | RPL | (23) | Allele-specific PCR | (65) |
| *ACE* | rs1799752‡ | D/I | - | 2.06<br>(1.46,2.91) | NA | 740/329 | RPL | (31) | Allele-specific PCR | (66) |
| *Folate metabolism* |  |  |  |  |  |  |  |  |  |  |
| *MTHFR* | rs1801133 | A/G | 0.36 | 1.25<br>(0.93,1.67) | 0.14 | 1830/3037 | RPL | (47) | RFLP | (67) |
| *Inflammatory response* |  |  |  |  |  |  |  |  |  |  |
| *IL6* | 14718 | C/G | - | 1.214<br>(0.88,1.67) | 0.24 | 230/188 | RPL | (51) | RFLP | (68) |
| *IL8* | rs2227306 | C/T | 0.61 | - | - | - | - | - | RFLP | (69) |
| *IL10* | rs1800896 | C/T | 0.45 | 1.27<br>(0.95,1.70) | > 0.05 | 635/571 | RPL | (50) | RFLP | (70) |
| *IL10* | rs1800872 | T/G | 0.76 | 3.01<br>(1.92,4.72) | < 10<sup>-4</sup> | 342/123 | Early PL | (49) | RFLP | (71) |

Legend: † Indel; ‡ Tandem repeat; EA/NEA - Effect allele/Non-effect allele; \*EAF (1000 G) – 1000 Genomes project effect allele frequency in Europeans

**Table 2.** Associations between 12 previously reported genetic variants and RPL in the REPLIK study.

| Locus name | Chr:Position | Variant ID | EA/NEA | EAF<br>cases/controls | N cases / N<br>controls | EAE | OR (95% CI) | P-value |
| --- | --- | --- | --- | --- | --- | --- | --- | --- |
| *F2* | 11:46761055 | rs1799963 | A/G | 0.011/0.72 | 110/46 | 0.49 | 1.64<br>(0.23, 32.48) | 0.66 |
| *F5* | 1:169519049 | rs6025 | T/C | 0.014/0.71 | 114/46 | -0.22 | 0.80<br>(0.15, 5.92) | 0.80 |
| *F7* | 13:113773159 | rs6046 | G/A | 0.48/0.070 | 75/46 | 0.34 | 1.40<br>(0.65, 3.04) | 0.38 |
| *GP1A* | 5:52347369 | rs1126643 | T/C | 0.35/0.23 | 81/46 | 0.35 | 1.43<br>(0.85,2.44) | 0.18 |
| *ESR1* | 6:152163335 | rs2234693 | T/C | 0.54/0.48 | 110/106 | -0.18 | 0.84<br>(0.57, 1.23) | 0.37 |
| *ADRB2* | 5:148206473 | rs1042714 | G/C | 0.33/0.25 | 81/46 | 0.38 | 1.47<br>(0.86,2.56) | 0.16 |
| *ENOS* | 15:35147732-35262040 | Intron-4†<br>VNTR | B/A | 0.55/0.12 | 102/46 | 0.15 | 1.16<br>(0.63,2.08) | 0.63 |
| *ACE* | 17:61565890 | rs1799752‡ | D/I | 0.35/0.0023 | 114/46 | -0.048 | 0.95<br>(0.59, 1.54) | 0.85 |
| *MTHFR* | 1:11856378 | rs1801133 | A/G | 0.21/0.52 | 114/46 | 0.33 | 1.40<br>(0.80, 2.51) | 0.25 |
| *IL6* | 7:22766840 | 14718 | C/G | 0.55/0.41 | 106/86 | 0.11 | 1.10<br>(0.77,1.64) | 0.57 |
| *IL8* | 4:74607055 | rs2227306 | C/T | 0.39/0.61 | 114/106 | 0.11 | 1.10<br>(0.76,1.64) | 0.60 |
| *IL10* | 1:206946897 | rs1800896 | C/T | 0.54/0.46 | 114/106 | 0.47 | 1.60<br>(1.07, 2.42) | 0.025 |
| **GRS** |  |  |  |  | **114/106** |  | **1.56**<br>**(1.21,2.04)** | **8.7x10<sup>-4</sup>** |

2 Legend: † Indel; ‡ Tandem repeat; EA/NEA: Risk allele/Alternate allele; EAF: Effect allele frequency; EAE: Estimated allelic effect (beta); GRS, genetic risk 3 score.

### Association analysis

We performed single-variant association analyses between each of genotyped variants and RPL. We used logistic regression for single-variant analyses assuming a log-additive model of association, similarly to standard assumptions in GWAS. We report estimates of ORs along with their 95% confidence intervals (CIs).

### Genetic risk score calculation

We calculated the genetic risk score (GRS) using information from a set of 12 variants previously associated with recurrent pregnancy loss (RPL) (**Table 1**). Among genotyped variants, there are ten SNPs, rs1799752 at *ACE* gene is an insertion/deletion (indel), and another is a tandem repeat (VNTR) in the intron 4 of *ENOS* gene. The risk allele count for each SNP was weighted by its established effect size using previously published findings. The effect sizes were estimated from the odds ratios (ORs) in the form of beta coefficients (log(OR)) for association between GRS and RPL assuming an additive genetic model for association. The weighted GRS was corrected for missing genotypes by multiplying the score with the total number of variants and then dividing by the number of genotyped variants per person, i.e. the scores for people with less genotyped variants got more weight (54). The effect estimate for rs1800896 located at *IL10* was from a multi-ethnic study, discordant from our study’s ethnic descent. We therefore used the present study effect estimate, which was still smaller than that published for the other *IL10* variant. In addition, for the variants in *F7, GP1A, ADRB2* and *IL8* there were no published studies for their association with RPL. Hence, we used the effect sizes from our data as weights. As a sensitivity check we also calculated the GRS using weights from our study only. Finally, we calculated an unweighted GRS and evaluated its effect on RPL.

### Receiver operating characteristic analysis

Next, we performed receiver operating characteristic (ROC) analysis to determine the predictive value of the estimated GRS in RPL. The efficacy of the GRS prediction is measured using the area under the curve (AUC) which is the statistic calculated on the observed case scale. The statistical analyses were conducted using the statistical software pROC package in R along with other functions (55).

## Results

### Association analysis

We tested the 13 genetic variants for association with RPL in the REPLIK study from Ukraine. Within this variant set only rs1800896 at *IL10* gene was nominally associated with RPL (OR[95% CI]: 1.60[1.07, 2.42], *P*=0.025, **Table 2**). Whereas the other variants did not reach nominal significance, five out the eight with an effect estimate available from literature were in the same direction of the effect in our study as in the published ones (**Table 1, Table 2**). The sample sizes varied greatly for the genotyped SNPs, whereas the GRS accounted for the variable number of genotyped SNPs per individual. The association between RPL and the GRS, based on published effect estimates, revealed a statistically significant association (*P=*8.7×10^−4^) in the REPLIK study. The combined effect of all tested variants resulted in a 1.56 times (95% CI [1.21, 2.04]) increased odds of RPL between cases and controls. The sensitivity analysis using the GRS with weights from our study effect estimates resulted in an OR of 1.83 (95% CI [1.34, 2.57], *P*=3.0×10^−4^). The unweighted GRS was also associated with an increased risk of RPL with an OR of 1.16 (95% CI [1.05, 1.29], *P*=2.0×10^−3^).

### Receiver Operator Characteristic analysis

The ROC analysis showed an area under the curve (AUC) of 0.64 (95% CI [0.57, 0.72]). This indicates a moderate to high ability for the GRS to correctly classify women with and without RPL. The sensitivity of 72% at the best discriminating point implies that the GRS can effectively identify women having experienced pregnancy losses (**Figure 1**). The AUCs for the GRS using our study effect estimates as weights and for the unweighted GRS were 0.64 (95% CI [0.57, 0.71]) and 0.62 (95% CI [0.55, 0.70]), respectively.

**Figure 1.**
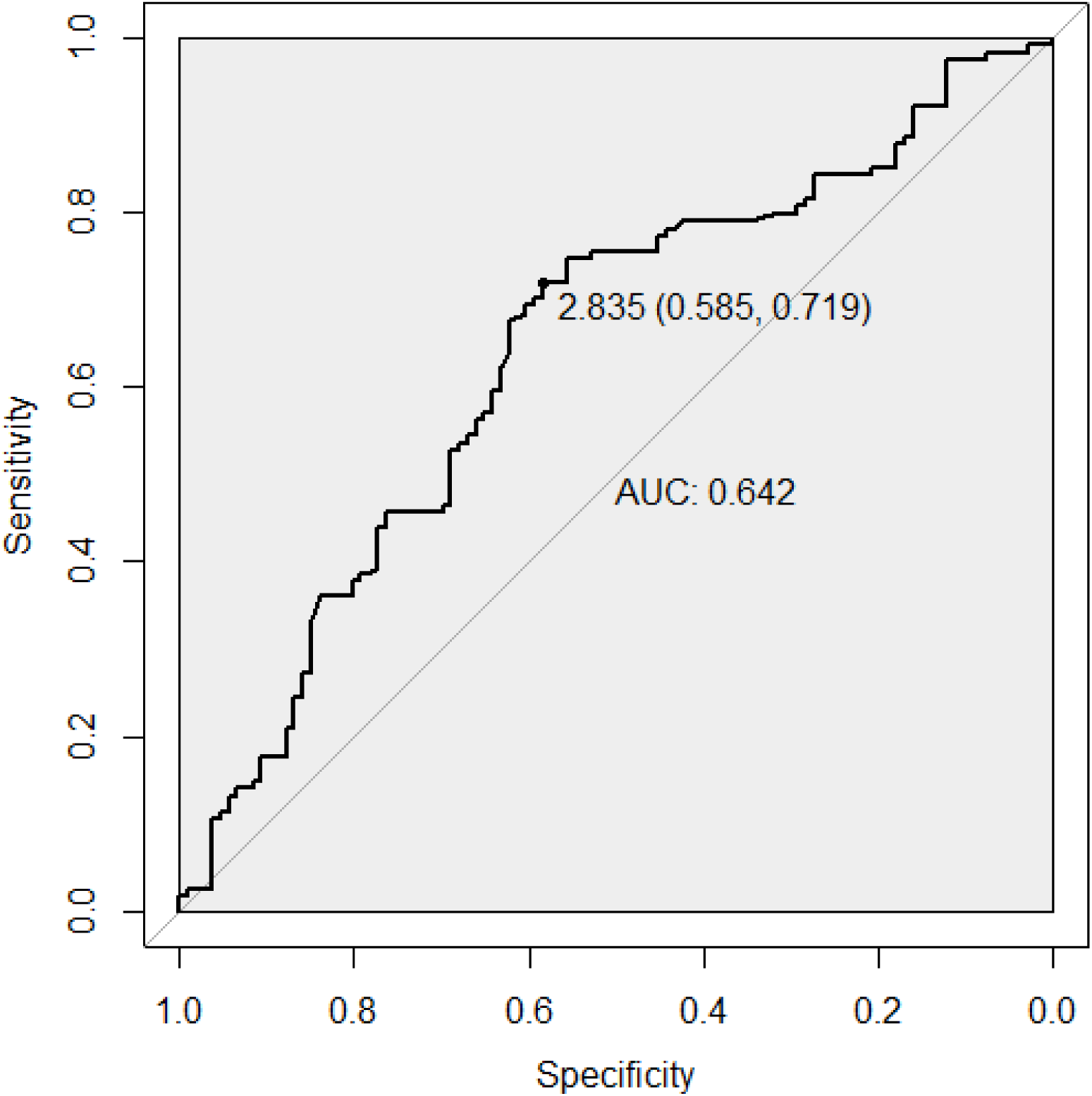
Receiver operator charateristic (ROC) curve for the predictive ability of the genetic risk score (GRS) for recurrent pregnancy loss. The best predictive point is shown with the ideal cut-off for the GRS and with estimates for specificity and sensitivity at that point. *Abbreviation: AUC, area under the curve.*

## Discussion

In this study we evaluated the combined effect of 12 genetic variants on RPL risk through GRS implemented in a case-control REPLIK study from the Ukraine. We showed that even with such a small number as 12 variants, when carefully chosen, we can already achieve predictive ability using weighted GRSs.

For some of the variants used (within *F2, F5, ACE, IL10*) we had previous evidence for their association with RPL (17,31,49), whereas others (within *F7, GP1A, ESR1, ADRB2, ENOS, MTHFR, IL6, IL8*) were chosen for their hypothesised biological mechanisms (18,19,23,27,28,47,51,52,56). The difficulty in establishing genetic associations with the complex condition of RPL was reflected in our single-variant analyses, i.e. we could not confirm the associations with RPL for majority of the variants with our available sample. A likely contributing factor was the relatively small sample size for some of the variants due to our genotyping strategy. For the *IL10* variant, having the whole sample genotyped, we demonstrated its association with RPL with a similar to previous study effect estimate (49).

It has been acknowledged that the discrete genetic variants for RPL have a relatively low sensitivity and specificity (3). Each individual case of idiopathic RPL usually cannot be explained by one risk factor and should be treated as a multifactorial condition (9). Indeed, GWAS for complex traits have shown that individual genetic variants usually provide relatively modest contribution to the trait variability in terms of their per-allele effect size, typically in the per-allele effects being within the range of 5-10% increase in risk in relation to that of risk estimated in general population (57), and hence require large sample sizes for detection. Therefore, it may be more effective to evaluate the risk of RPL using a panel of population-specific low-effect genetic markers, representing distinct physiological gene networks. Our selection of the gene panel based on the hypothesised biological pathways proved successful since by combining their effects, we could already predict the risk of RPL in our cohort. The results were not influenced by the selection of the weights, as our sensitivity analyses showed. It is worth noticing that we achieved an AUC of 0.64 (95% CI 0.57-0.72) with already 12 SNPs. A recent study combining millions of SNPs into genome-wide polygenic scores for several complex diseases achieved AUCs of similar strength. For example, an AUC of 0.63 was reported for inflammatory bowel disease from a GRS consisting of 6.9 million SNPs (58). Taken together, our results are important considering that RPL is a laborious and expensive phenotype to collect. Since the start of large-scale genome-wide association studies, RPL has lacked novel insights establishing its underlying genetic mechanisms with no major publications probably due to clinical requirements to the phenotype definition. However, our investigation suggests that even a small number of SNPs in appropriately defined cases and controls can be used for predictive purposes.

One important limitation of RPL genetic association studies arises from the fact that even though the differences in population frequencies of studied genetic polymorphisms may be significant, ethnicity-specific associations are rarely addressed. Moreover, a vast majority of studies are performed on European and American Caucasians, as well as, less frequently, Southern or Eastern Asians (5,23,59). On one hand, this situation allows increasing the significance of discovered effects in European populations through meta-analyses (17). On the other hand, it complicates adoption of such effect estimates for other ethnic groups due to possible difference in allele frequencies and effect size estimates between populations and ethnic groups (23,56).

In summary, with the careful selection of the DNA variant set and the implementation of methods such as the GRS, we can predict susceptibility to complex multifactorial conditions such as RPL. With the hope of future well-powered studies, especially GWAS, adding to the knowledge of biological pathways not hypothesised previously for RPL, we will able to further improve the prediction and identification of those at risk for RPL.

## Acknowledgements

IP is funded in part by the World Cancer Research Fund (WCRF UK) and World Cancer Research Fund International (2017/1641), the Wellcome Trust (WT205915).

